# Nucleolin loss-of-function leads to aberrant FGF signaling and craniofacial anomalies

**DOI:** 10.1101/2021.09.14.460382

**Authors:** Soma Dash, Paul A. Trainor

**Affiliations:** Stowers Institute for Medical Research, Kansas City, MO, USA; Department of Anatomy and Cell Biology, University of Kansas Medical Center, Kansas City, KS, USA

## Abstract

rRNA transcription and ribosome biogenesis are global processes required for growth and proliferation of all cells, yet perturbation of these processes in vertebrates leads to tissue-specific defects termed ribosomopathies. Mutations in rRNA transcription and processing proteins often lead to craniofacial anomalies, however the cellular and molecular reasons for this are poorly understood. Therefore, we examined the function of the most abundant nucleolar phosphoprotein, Nucleolin (Ncl), in vertebrate development. We discovered that Nucleolin is dynamically expressed during embryonic development with high enrichment in the craniofacial tissues. Consistent with this pattern of expression, *ncl* homozygous mutant (*ncl^-/-^*) zebrafish present with craniofacial anomalies such as mandibulofacial hypoplasia. We observe that *ncl^-/-^* mutants exhibit decreased rRNA synthesis and p53-dependent neuroepithelial cell death. In addition, the half-life of *fgf8a* mRNA is reduced in *ncl^-/-^* mutants, which perturbs Fgf signaling, resulting in misregulation of Sox9a mediated chondrogenesis and Runx2 mediated osteogenesis. Exogenous addition of human recombinant FGF8 to the mutant zebrafish significantly rescues the cranioskeletal phenotype, suggesting that Nucleolin regulates osteochondroprogenitor differentiation during craniofacial development by post-transcriptionally regulating Fgf signaling. Our work has therefore uncovered a novel tissue-specific function for Nucleolin in rRNA transcription and growth factor signaling during embryonic craniofacial development.

## Introduction

The craniofacial complex consists of the primary sense organs, central and peripheral nervous systems, and musculoskeletal components of the head and neck. Craniofacial development is an intricate process that involves coordinated interaction of all three germ layers and is sensitive to environmental and genetic insults resulting in craniofacial disorders. Despite advances in sequencing, many affected individuals have an unknown genetic diagnosis. Therefore, it is necessary to identify novel genetic factors and understand the cellular and molecular mechanisms that regulate normal craniofacial development, which may also aid in identifying potential therapeutic targets to prevent and/or ameliorate craniofacial diseases.

Ribosome biogenesis is essential for cell growth and survival because ribosome quantity and quality dictate the translation of mRNA into proteins. Transcription of ribosomal DNA (rDNA) by RNA Polymerase (Pol) I in the nucleolus generates a 47S pre-ribosomal RNA (pre-rRNA). The 47S pre-rRNA is then cleaved and processed into 18S, 5.8S and 28S rRNAs. These rRNAs together with Pol III transcribed 5S rRNA associate with ribosomal proteins and accessory proteins to form ribosomes^1^. The transcription and processing of pre-rRNA requires RNA Pol I together with associated proteins such as UBTF and SL-1^2,3^ as well as rRNA processing proteins including Tcof1^4^, Nol11^5^, Wrd43^6^ and Fibrillarin^7^. Interestingly, when Pol I subunits or any of the associated factors are disrupted or mutated in zebrafish, xenopus or mice, it results in developmental defects that mostly affect craniofacial cartilage and bone differentiation^6,8–12^. This raises the question of why disruptions in these ubiquitously expressed genes, which are required in a global process, result in tissue specific craniofacial anomalies. One hypothesis is that neural crest cells (NCC), which are the progenitors of most of the craniofacial bone and cartilage, are more proliferative or metabolically active than non-neural crest cells. In addition, NCCs undergo major cytoskeletal changes during their epithelial-to-mesenchymal transition, which requires high levels of new protein synthesis and hence, more rRNA transcription. Another hypothesis is that RNA Pol I subunit, associated proteins and rRNA processing proteins have other non-ribosome functions, which together with the regulation of rRNA synthesis make the craniofacial skeleton more susceptible to disruption.

Nucleolin is a major nucleolar protein and rRNA processing protein as well as an mRNA and DNA binding protein^13,14^. Among the top ten highly enriched genes in neural crest cells, *Ncl* is the only one involved in rRNA transcription^15^. Therefore, in this study we explore the hypothesis that Nucleolin regulates rRNA transcription as well as craniofacial specific gene expression and function during embryogenesis. We show that Nucleolin is essential for embryo survival and is required for craniofacial bone and cartilage development. Nucleolin regulates rRNA transcriptionally, *fgf8a* mRNA post-transcriptionally and p53 protein post-translationally. Consistent with this model, we demonstrate that exogenous human recombinant FGF8 can ameliorate the cranioskeletal defects as well as recover rRNA transcription in *ncl^-/-^* mutant embryos. Our work, therefore, has uncovered novel tissue-specific functions for Nucleolin in craniofacial development through regulating rRNA transcription and FGF signaling during embryogenesis.

## Results

### Nucleolin is dynamically expressed during craniofacial development

To understand the function of Nucleolin in vertebrate development, we characterized its expression during zebrafish embryogenesis (Fig. 1). Nucleolin is maternally expressed at 1.5 hour post fertilization (hpf) and remains ubiquitously expressed through gastrulation (3 hpf), early neurulation (12 hpf) and axial segmentation (18 hpf) (Fig. 1A-D). At 24 hpf, the expression of Nucleolin is still ubiquitous, however it is enriched in the eye and midbrain-hindbrain boundary (MHB) (Fig. 1E). In 36 and 72 hpf zebrafish embryos, elevated expression of Nucleolin is observed within the eye, pharyngeal arches and brain (Fig. 1F,G). These expression analyses demonstrate that Nucleolin is dynamically expressed during embryogenesis with enriched expression in craniofacial tissues. Further, this suggests that Nucleolin may be required for proper craniofacial morphogenesis.

**Figure 1.**
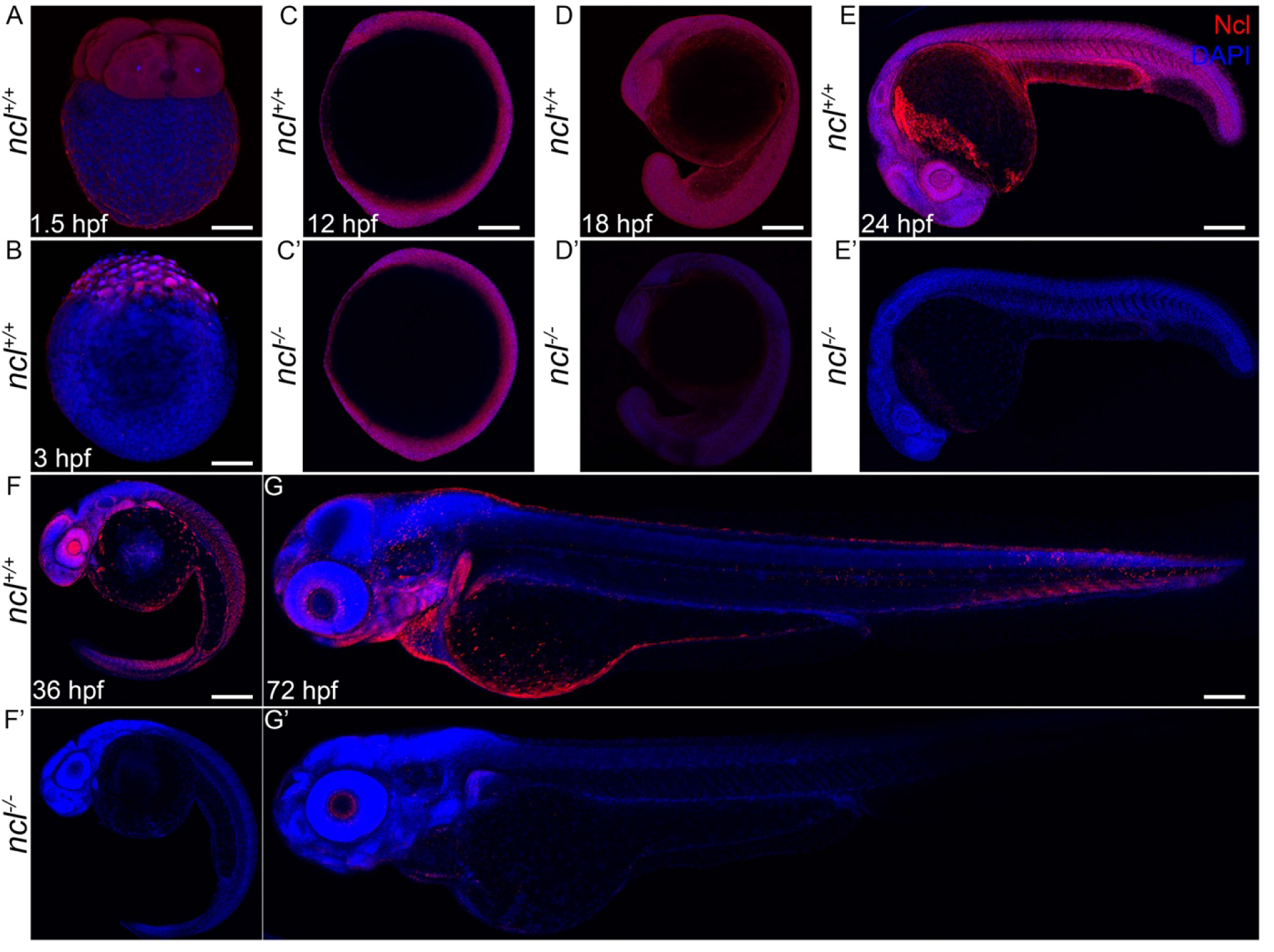
Ncl expression during zebrafish development. (A) During embryogenesis, Nucleolin is ubiquitously expressed in the cytoplasm of 4 cell stage wildtype embryo at 1.5hpf. (B) Similarly, 3hpf embryos also have ubiquitous cytoplasmic expression of Nucleolin. (C-C’) At 12hpf, *ncl^+/+^* and *ncl^-/-^* embryos exhibit similar Nucleolin expression in the nucleus and cytoplasm in most cells of the embryos. (D-D’) By 18hpf, the expression of Nucleolin in *ncl^+/+^* embryos is confined to the nucleus and is ubiquitous. In the *ncl^-/-^* embryos, the expression pattern of Nucleolin is similar to that of wildtype, however, the expression level is significantly lower than that of the wildtype. (E-E’) By 24 hpf, the expression of Nucleolin is still ubiquitous with higher levels in the eye and the midbrain-hindbrain boundary in *ncl^+/+^* embryos, while it is absent in *ncl^-/-^* embryos. (F) At 36 hpf, the expression of Nucleolin becomes specific to the craniofacial region in the pharyngeal arches as well as the eye. (G) In 3 dpf wildtype zebrafish, Nucleolin expression is specific to the jaw of the embryo. (F’-G’) In the *ncl^-/-^* mutants, there is no expression of Nucleolin. Scale bar denotes 35 μm for A-B, 70 μm for C-C’, 140 μm for D-D’, 250 μm for E-E’ and 300 μm for F-G’.

### Mutations in zebrafish *ncl* result in craniofacial anomalies

To test our hypothesis that Nucleolin functions during craniofacial development, we characterized the phenotype of the *ncl* mutant zebrafish line, *ncl^hi2078Tg^*. *ncl^hi2078Tg^* was generated by insertional mutagenesis of exon 1, which disrupts *ncl* transcription^16^, dramatically reducing the levels of Nucleolin protein between 18 hpf and 24 hpf (Fig 1C’-G’), thereby establishing *ncl^hi2078Tg^* zebrafish as *ncl^-/-^* mutants.

*ncl^-/-^* embryos are phenotypically distinguishable from their WT siblings at 24 hpf by the presence of necrosis in the craniofacial region (Fig. 2A-B). At 36hpf, *ncl^-/-^* embryos have a beak-like frontonasal prominence and misshapen midbrain-hindbrain boundary (MHB) (Fig. 2C-D’). By 3 days post fertilization (dpf), *ncl^-/-^* mutants display craniofacial anomalies such as mandibular hypoplasia in addition to a misshapen MHB (Fig. 2E-F’). 5dpf *ncl^-/-^* embryos continue to exhibit craniofacial anomalies and fail to inflate their swim bladders (Fig. S1A-A’), which collectively leads to their lethality between 6-10 dpf (Fig. S1 D-D’).

**Figure 2.**
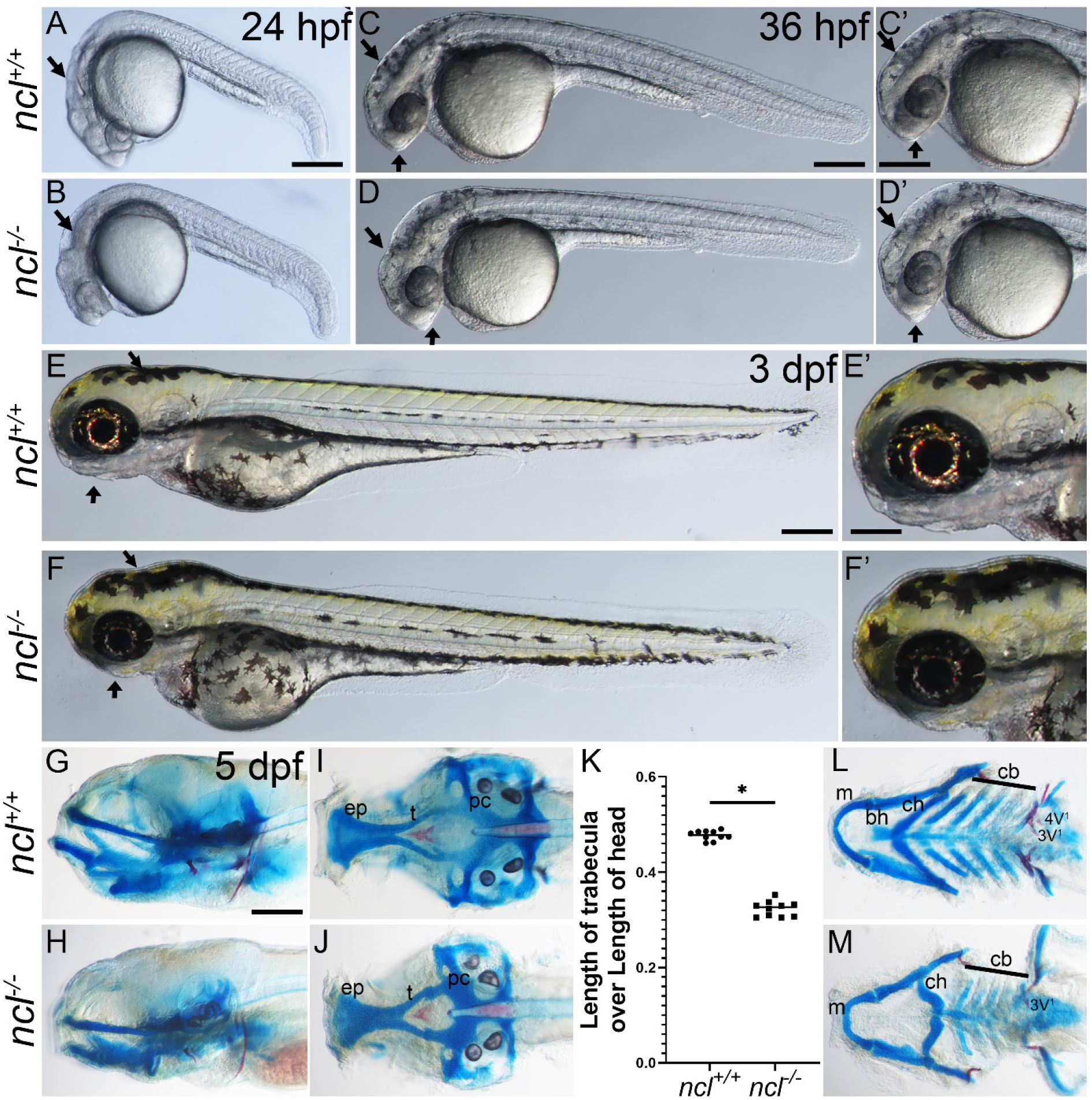
*ncl^-/-^* mutants exhibit craniofacial defects. (A-B) Compared to 24 hpf *ncl^+/+^* clutch mates, *ncl^-/-^* mutants have necrotic tissue (indicated by black arrows) in the craniofacial region. (C-D) By 36 hpf, the frontonasal prominence of *ncl^-/-^* mutants has a beak like appearance and the midbrain-hindbrain boundary is misshapen compared to *ncl^+/+^* siblings (indicated by black arrows). The craniofacial region is magnified in C’-D’. (E-F) At 3 dpf, the *ncl^-/-^* mutants have smaller jaws and a misshapen head (indicated by black arrows). The craniofacial region is magnified in E’-F’. (G-H) Skeletal preparations of 5 dpf wildtypes and *ncl^-/-^* mutants reveal defects in the cartilages of the jaw. (I-J) In the neurocranium, the chondrocytes in the ethmoid plate (ep) are delayed in development, and the trabeculae are smaller and wider compared to the wildtype zebrafish at the same stage. (K) Quantification of the length of trabecula/ presphenoid in *ncl^+/+^* and *ncl^-/-^* embryos as a ratio of the length of the head measured from anterior most point of ethmoid plate to the posterior most point of parachordal (pc). (L-M) In the viscerocranium, Meckel’s cartilage is bent, the basihyal is missing, the polarity of the ceratohyal is inverted and the ceratobranchials are hypoplastic. In addition, the mutants have hypoplastic teeth and the 4V^1^ teeth are missing. Abbr. ep: ethmoid plate, t: trabecula, pc: parachordal, m: meckel’s, bh: basihyal, ch: ceratohyal, cb: ceratobranchial. Scale bars denotes 200 μm for A-F, 50 μm for C’-F’, 70 μm for G-J, L-M.

To further characterize the craniofacial defects in *ncl^-/-^* embryos, we performed skeletal staining with Alcian blue and Alizarin red for cartilage and bone, respectively. At 3 dpf, *ncl^-/-^* embryos display hypoplastic Meckel’s and ceratohyal cartilages compared to their *ncl^+/+^* siblings. (Fig. S1B-B’). In 5 dpf *ncl^-/-^* embryos, the craniofacial cartilages are severely hypoplastic (Fig. 2G-M). In the neurocranium, the trabecula and parasphenoid, which forms the base of the skull^17^, are smaller in *ncl^-/-^* mutants compared to *ncl^+/+^* siblings (Fig. 2I-J). In *ncl^+/+^* larvae, both edge and medial chondrocyte populations are present and are demarcated by differential Alcian blue staining. However, in *ncl^-/-^* larvae, cells in the ethmoid plate exhibit uniform Alcian blue staining, suggesting that differentiation of medial chondrocytes in the ethmoid plate may be perturbed. In addition, the trabecula and parasphenoid are smaller in *ncl^-/-^* larvae compared to *ncl^+/+^* larvae (Fig. 2K). In the viscerocranium, Meckel’s cartilage is misshapen and the ceratohyal exhibits reverse polarity while the basihyal is missing in *ncl^-/-^* larvae. Furthermore, the posterior pharyngeal arch derived ceratobranchials are hypoplastic (Fig. 2L-M) and osteogenesis of the teeth is incomplete in *ncl^-/-^* larvae. At 8 dpf and 10 dpf, the polarity of the ceratohyal is similar between *ncl^+/+^* and *ncl^-/-^* larvae, suggesting that *ncl^-/-^* larvae may have delayed cranioskeletal development. However, Meckel’s cartilage remains misshapened and the ceratobranchials remain hypoplastic in 8 dpf *ncl^-/-^* mutant zebrafish (Fig. S1 B-C’ and E-F’). This establishes Nucleolin as being essential for proper cranioskeletal development, and *ncl^-/-^* mutant zebrafish as a new model for understanding the etiology and pathogenesis of craniofacial anomalies.

### *ncl^-/-^* zebrafish have diminished rRNA synthesis

Molecularly, Nucleolin binds to and modifies histones on the promoter of rDNA, and thereby regulates rRNA transcription, which is a rate-limiting step of ribosome biogenesis^18^. In addition, Nucleolin is a component of the U3 snoRNA, and thus binds to pre-rRNA to process and splice it^13,19,20^. Therefore, we hypothesized that *ncl* loss-of-function would lead to diminished rRNA transcription and processing.

To validate our hypothesis, we quantified rRNA transcription by assaying for the intergenic regions of the 47S pre-rRNA - 5’ETS, ITS1 and ITS2 as well as 18S rRNA by qRT-PCR. We observed a significant reduction of all four amplicons in *ncl^-/-^* embryos compared to *ncl^+/+^* siblings, suggesting that Nucleolin is necessary for rRNA transcription in zebrafish (Fig. 3A). We then performed RNA immunoprecipitation using a Nucleolin specific antibody and observed that Nucleolin binds to the 5’ETS and ITS1 but not to ITS2 and 18S regions of the 47S pre-RNA (Fig. 3B), consistent with the known function of Nucleolin *in vitro*^18^.

**Figure 3.**
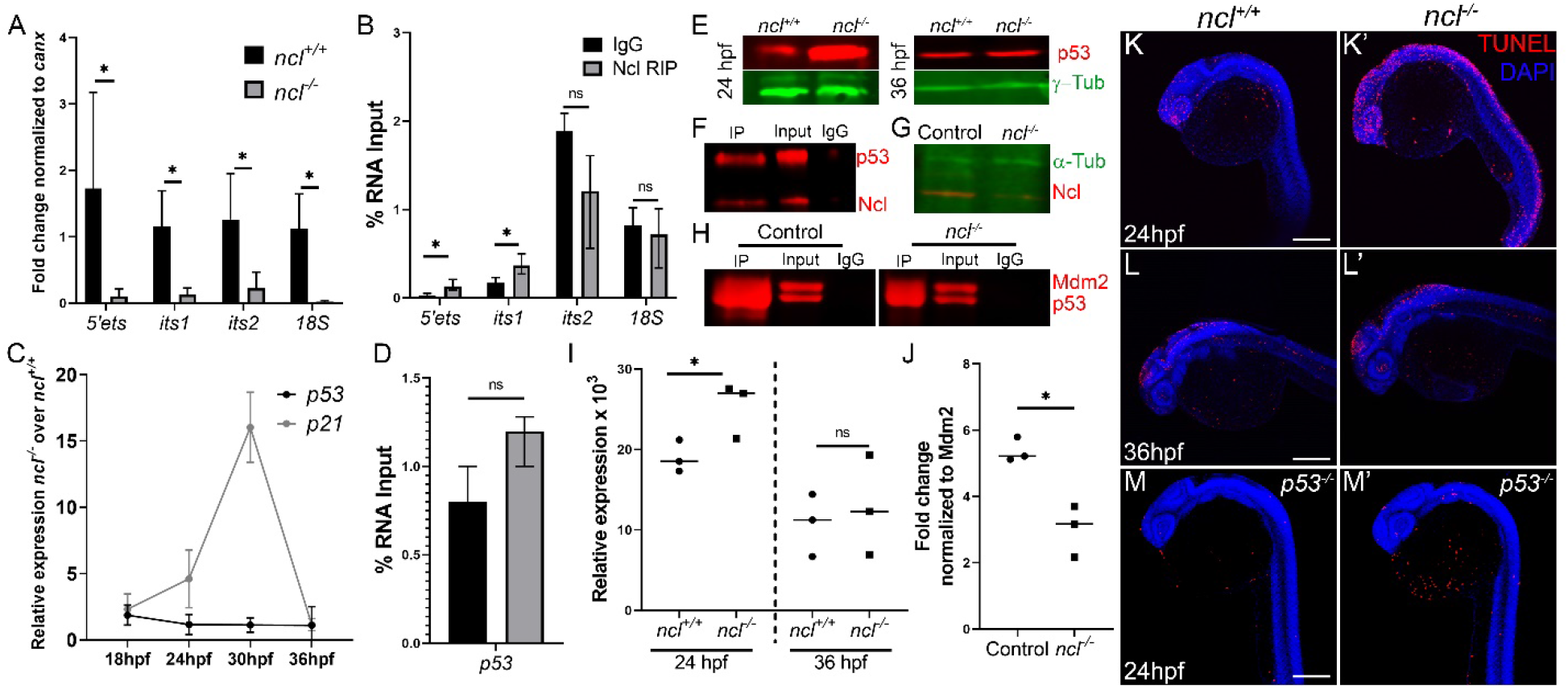
Nucleolin is required for rRNA transcription and p53 regulation. (A) qPCR for 5’ETS, ITS1, ITS2 and 18S segment of the pre-rRNA in *ncl^+/+^* and *ncl^-/-^* zebrafish indicates that rRNA transcripts are significantly lower in *ncl^-/-^* embryos compared to *ncl^+/+^* siblings. *canx* was used as an internal control. (B) RNA immunoprecipitation using a Nucleolin specific antibody indicates that Nucleolin binds to the 5’ETS and ITS1 region of the 47S rRNA but not to the ITS2 or 18S in wildtype zebrafish. (C) *p53* transcript expression is not significantly altered in *ncl^-/-^* mutant zebrafish between 18-36 hpf, however, its downstream target p21 is significantly higher between 24-30 hpf in the *ncl^-/-^* mutants compared to wildtype zebrafish. (D) Nucleolin and IgG binding to *p53* mRNA is similar in wildtype zebrafish as observed by RNA immunoprecipitation. (D) p53 protein levels are higher in *ncl^-/-^* mutants at 24hpf compared to *ncl^+/+^* sibling and comparable between *ncl^+/+^* and *ncl^-/-^* embryos at 36 hpf as observed by western blotting. γ-tubulin was used as a loading control. (F) Immunoprecipitation with a Nucleolin specific antibody followed by western blotting for p53 and Nucleolin indicates that p53 and Nucleolin bind to each other in wildtype zebrafish. (G) In *ncl^-/-^* mutants, Nucleolin expression is significantly reduced compared to controls. α-tubulin was used as a control. (H) At 28hpf, control zebrafish have higher binding of Mdm2 and p53 compared to mutant zebrafish. (I) Quantification of p53 protein levels in 24 hpf and 36 hpf *ncl^+/+^* and *ncl^-/-^* embryos. (J) Quantification of p53-Mdm2 binding in *ncl^+/+^* and *ncl^-/-^* embryos (K-K’) *ncl^-/-^* mutants have more TUNEL positive cells at 24 hpf compared to *ncl^+/+^* siblings. (L-L’) By 36 hpf, apoptosis is confined to the midbrain-hindbrain boundary in both *ncl^+/+^* and *ncl^-/-^* embryos. (M-M’) On a *p53^-/-^ mutant* background, the number of TUNEL positive cells in both *ncl^+/+^* and *ncl^-/-^* embryos at 24hpf is reduced. Scale bar denotes 70 μm for K-M’.

Decreased rRNA transcription has previously been shown *in vitro* to result in increased free ribosomal proteins in cells, which then interact with Mdm2, changing its conformation such that Mdm2 can no longer bind to p53 and ubiquitinate it for degradation. Consequently, p53 accumulates in the cell and results in p53 dependent cell death^21^. To test if this mechanism holds true in *ncl^-/-^* embryos, we first performed qPCR and western blot to assess for p53 expression. We observed that the *p53* transcript level is not significantly changed in *ncl^-/-^* embryos compared to *ncl^+/+^* siblings between 18 hpf and 36 hpf (Fig. 3C). In contrast, p53 protein levels are significantly upregulated at 24 hpf, however the difference between *ncl^+/+^* and *ncl^-/-^* mutants subsides by 36 hpf (Fig. 3E, I). The downstream target of p53, *p21* is initially upregulated between 24-30 hpf, but then is considerably downregulated by 36 hpf (Fig. 3C). Nucleolin regulates p53 expression by various mechanisms including increasing p53 mRNA and/or protein stability^22,23^. We observe that in wildtype zebrafish tissues, Nucleolin does not bind to zebrafish *p53* mRNA (Fig. 3D). Instead, Nucleolin binds to p53 protein (Fig. 3F). Prior to examining the binding efficiency of Mdm2 and p53 in control and *ncl^-/-^* embryos, we confirmed that Nucleolin is indeed absent in *ncl^-/-^* embryos at 28hpf (Fig. 3G). We then performed immunoprecipitation for Mdm2 in control and *ncl^-/-^* embryos and immunoblotted for both Mdm2 and p53. We observe decreased pulldown of Mdm2 and p53 in *ncl^-/-^* embryos compared to control embryos indicating that the initial accumulation of p53 is a result of reduced interactions between Mdm2 and p53 (Fig. 3H-J). The downregulation of p53 at later stages might be due to a lack of Nucleolin stabilizing p53 protein to increase its half-life.

### p53-dependent cell death is increased in *ncl* mutant embryos

To examine if increased p53 activation and accumulation results in cell death and therefore necrotic craniofacial tissue in the mutants (Fig. 2A), we performed TUNEL staining at 24 hpf and observed a general increase in apoptosis in *ncl^-/-^* embryos compared to *ncl^+/+^* controls (Fig. 3K-K’). However, by 36 hpf, TUNEL positive cells were restricted to the MHB in *ncl^-/-^* embryos (Fig. 3L-L’), consistent with a reduction in p53 protein levels.

p53 activation can lead to increased apoptosis as well as decreased proliferation that collectively result in tissue hypoplasia. Therefore, we performed immunostaining with the G2/M phase marker phospho-histone H3 (pHH3) and observed no significant change in pHH3+ cells in 24 hpf *ncl^-/-^* mutants compared to controls (Fig S2 A-B). Consistent with the decrease in p53 and p21 expression at 36 hpf, the number of pHH3 positive cells is significantly increased in *ncl^-/-^* embryos especially at the MHB at 48 hpf (Fig. S2 C,D,G). We also performed EdU staining on *ncl^+/+^* and *ncl^-/-^* zebrafish at 48 hpf to label cells in S-phase of the cell cycle and observed that the mutants have a higher number of EdU positive cells (Fig. S2 E,F,H). The increase in proliferation may explain how *ncl^-/-^* embryos survive beyond 24 hpf, even though they exhibit higher apoptosis.

We hypothesized that the increased apoptosis in *ncl^-/-^* embryos is p53 dependent and that genetically inhibiting p53 levels would suppress cell death and rescue the craniofacial anomalies in *ncl^-/-^* embryos. Therefore, we crossed the *tp53^M214K/M214K^* allele (hereafter referred to as *p53^-/-^*) into the background of *ncl^-/-^* mutant zebrafish to generate *ncl^-/-^;p53^-/-^* double mutants. Consistent with our hypothesis, TUNEL staining revealed a reduction in apoptotic cells in 24 hpf *ncl^-/-^;p53^-/-^* embryos compared to *ncl^-/-^* embryos (Fig 3L-L’). However, EdU labeling indicates that *ncl^-/-^;p53^-/-^* embryos have increased proliferation at 48 hpf similar to *ncl^-/-^* (Fig. S2 I-J), suggesting that the proliferation defects are not p53 dependent.

Removal of both the copies of *p53* and knocking down p53 using morpholinos altered jaw morphology in *ncl^-/-^* embryos (Fig. S3). Although the ceratohyal remained smaller compared to control siblings, its polarity was restored to normal. The basihyal, which was absent in *ncl^-/-^* mutant embryos is formed in *ncl^-/-^;p53^-/-^* embryos. However, chondrogenesis of the ceratobranchials was not improved. In addition, chondrogenesis in the neurocranium was disrupted. *ncl^-/-^; p53^-/-^* mutant zebrafish died around 10-12 dpf due to these cranioskeletal defects and a failure to inflate their swim bladders which is similar to *ncl^-/-^* mutant zebrafish. This suggests that while p53 accumulation results in apoptosis during early development, the skeletal defects in *ncl^-/-^* mutant zebrafish are not p53 dependent.

### NCC development is unaffected in *ncl^-/-^* mutant embryos

In vertebrates, NCC differentiate into most of the bones and cartilages of the craniofacial skeleton. To determine if defects in NCC induction and migration underlie the cranioskeletal malformations in *ncl^-/-^* mutants, we labeled NCC by crossing *sox10:egfp* transgenic zebrafish into *ncl^-/-^* mutant zebrafish. In addition, we immunostained *ncl^+/+^* and *ncl^-/-^* embryos with Zn-8, which labels the endodermal pouches to demarcate individual pharyngeal arches^24,25^. *sox10:egfp* labeling of pre- and post-migratory NCC and volumetric rendering of the pharyngeal arches revealed no significant change in the size of the pharyngeal arches in *ncl^-/-^* mutant embryos compared to *ncl^+/+^* embryos (Fig. S4 A-B). Comparison of *sox10:egfp* fluorescence intensity in the pharyngeal arches of *ncl^+/+^* and *ncl^-/-^* embryos suggests that NCC induction and migration to the arches are unaffected in the *ncl^-/-^* mutant embryos (Fig. S4 C). However, the pineal gland as well as the nasopharyngeal regions of *ncl^-/-^* mutants have reduced *sox10:egfp* expression suggesting that while NCC migration into the arches is unaffected, migration to other cranial regions may be altered. We further validated *sox10;gfp* transgene expression data by assessing endogenous Sox10 protein expression in *ncl^+/+^* and *ncl^-/-^* embryos. The number of Sox10+ cells in the pharyngeal region is similar between 36 hpf *ncl^+/+^* and *ncl^-/-^* mutant embryos (Fig. S4 D), confirming that NCC induction and migration into the pharyngeal arches occurs normally in *ncl^-/-^* mutants. We observed no alteration in the formation or segregation of the endodermal pouches in 36 hpf *ncl^-/-^* mutants as evidenced by a normal pattern of Zn-8 expression (Fig. S4 A). This implied that the osteochondrogenic anomalies in *ncl^-/-^* mutant embryos were not the result of abnormal pharyngeal pouch development.

### Chondrogenic and Osteogenic defects in *ncl^-/-^* mutants

Given the chondrogenic defects observed in *ncl^-/-^* mutant embryos, we next tested for altered chondrogenic markers in *ncl^+/+^* and *ncl^-/-^* embryos. Sox9a is a transcriptional factor that promotes NCC differentiation into chondrocytes^26^. *sox9a* mRNA and its downstream target, *col2a1a* are reduced in *ncl^-/-^* mutant embryos (Fig. 4A). The overall expression of Sox9a protein is also reduced in *ncl^-/-^* mutants, but significantly in pharyngeal arches 2-5, which differentiate into the ceratohyal and ceratobranchial cartilages (Fig. 4B). This provides a molecular explanation for the cartilage hypoplasia in *ncl^-/-^* mutant embryos.

**Figure 4.**
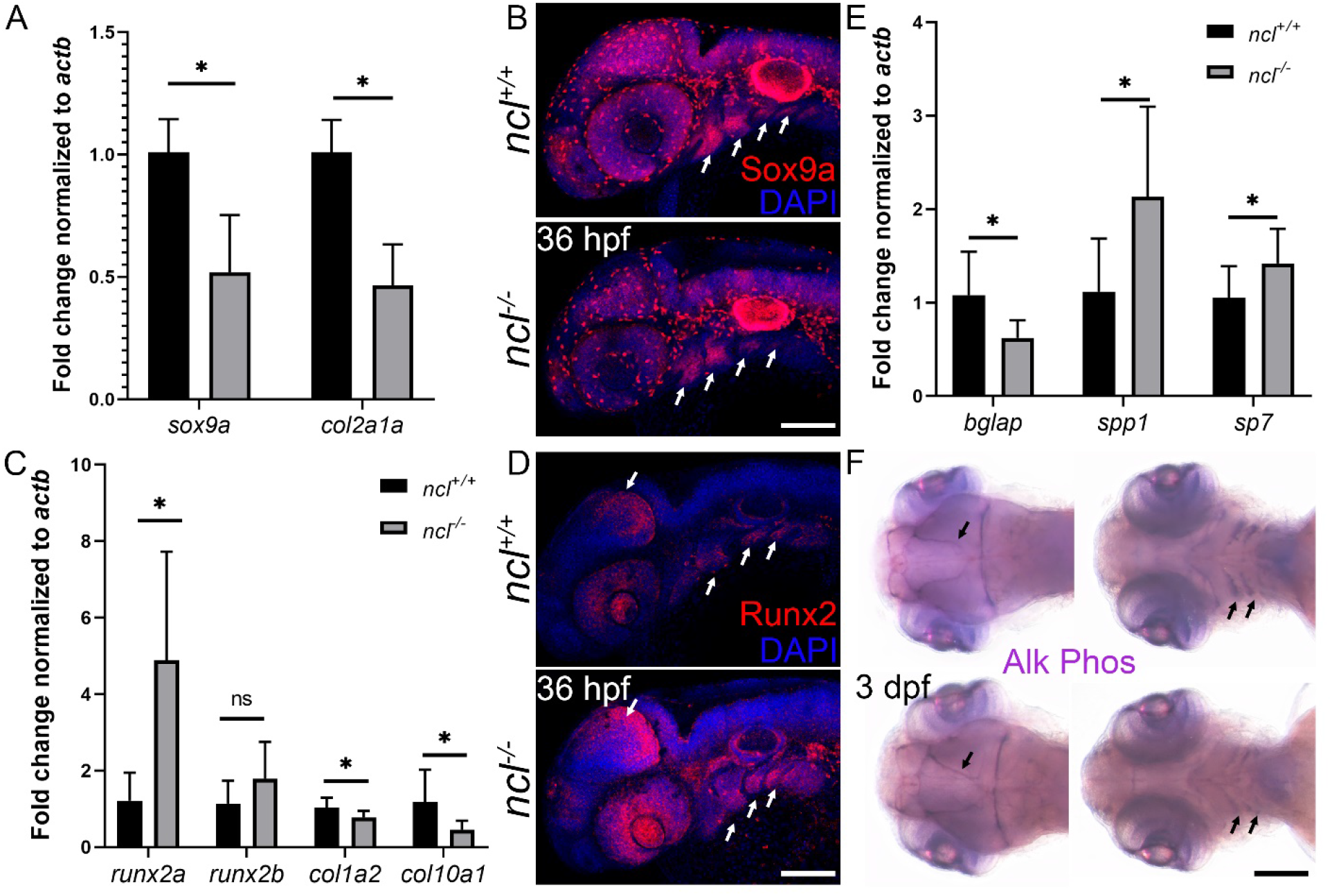
Chondrogenesis and osteogenesis defects in *ncl^-/-^* embryos. (A) qPCR revealed a significant downregulation of *sox9a* and *col2a1* chondrogenesis markers in 36 hpf *ncl^-/-^* embryos compared to *ncl^+/+^* embryos. *actb* was used as housekeeping control. (B) Sox9a protein expression is significantly reduced in branchial arches 2-5 (white arrows) in *ncl^-/-^* embryos at 36 hpf. (C) qPCR of osteogenesis markers in 36 hpf *ncl^+/+^* and *ncl^-/-^* embryos indicates significant upregulation in *runx2a* transcript and downregulation in both *col1a2* and *col10a1* in *ncl^-/-^* embryos. (D) Runx2 protein expression is significantly increased in the midbrain-hindbrain boundary and branchial arches 2-5 (white arrows) in *ncl^-/-^* embryos at 36 hpf. (E) qPCR indicates a significant upregulation of early osteoblast markers, *bglap* and *spp1,* and downregulation of the late osteoblast marker *sp7* in 36 hpf *ncl^-/-^* embryos compared to controls (F) Alkaline phosphatase staining of 3 dpf *ncl^+/+^* and *ncl^-/-^* embryos reveals the distorted shape of the cranial sutures (dorsal view) and reduced staining in the lower jaw (ventral view, black arrows) in ventral view (indicated by black arrows). Scale bar denotes 100 μm for B, D and F.

To determine the basis for the osteogenic defects in *ncl^-/-^* mutant embryos, we assessed for altered osteogenic markers in *ncl^+/+^* and *ncl^-/-^* embryos. Runx2a and Runx2b are transcription factors that regulate NCC differentiation to bone^27^. *runx2a* is upregulated in the *ncl^-/-^* mutant embryos, as is *runx2b*, but *runx2b* upregulation is not statistically significant (Fig. 4C). We examined the expression of Runx2a and Runx2b in *ncl^-/-^* and *ncl^+/+^* embryos using a pan-Runx2 antibody, which revealed that Runx2 protein is upregulated in craniofacial regions especially in the MHB and pharyngeal arches (Fig. 4D). Runx2 overexpression has been previously observed to result in premature osteoblast differentiation^28^, which leads to a reduced pool of osteoblasts. In agreement with this finding, the expression of Runx2a and Runx2b downstream targets, *col1a2* and *col10a1* are reduced in *ncl^-/-^* embryos compared to *ncl^+/+^* embryos. Furthermore, we observe an increase in early osteoblast markers, *spp1* (osteopontin) and *sp7* and a concomitant decrease in late osteoblast marker, *bglap* (osteocalcin) in 36 hpf *ncl^-/-^* mutant embryos. To confirm the presence of prematurely differentiated osteoblasts, we stained *ncl^+/+^* and *ncl^-/-^* mutant embryos for alkaline phosphatase activity, which is endogenously high in primary osteoblasts^29^. At 3 dpf (Fig. 4F) and 5 dpf (Fig. S5), alkaline phosphatase staining is significantly diminished in the lower jaw of *ncl^-/-^* embryos compared to *ncl^+/+^* embryos. Interestingly, in the neurocranium, osteoblasts in the cranial sutures at both 3 dpf and 5 dpf stain for alkaline phosphatase, which reveals the shape of the sutures in *ncl^-/-^* mutant embryos is altered, consistent with the effects of Runx2 mutations in humans^30,31^.

### Fgf8a expression is reduced in *ncl^-/-^* mutant embryos

In mice, Fgf8 signaling upregulates chondrogenic genes such as *Sox9* and *Col2a1* while inhibiting osteogenic genes including *Runx2*^32,33^. Similarly, Fgf signaling regulates *Sox9a* in zebrafish^34^. Interestingly, *fgf8a* mutant zebrafish have craniofacial defects comparable to *ncl^-/-^* embryos, especially in the cartilages of the viscerocranium^35^, suggesting a potential link between Nucleolin and Fgf signaling during chondrogenesis. Therefore, we examined the expression of Fgf8a in *ncl^+/+^* and *ncl^-/-^* embryos and observed a significant general downregulation of Fgf8a in *ncl^-/-^* mutant embryos (Fig. 5A). We examined the expression of *fgf8a* mRNA by qPCR at four stages between 18-36 hpf in *ncl^-/-^* mutant embryos and discovered that as the expression of *ncl* decreases in the mutants, the expression of *fgf8a* also declines (Fig. 5B). In silico analysis of *fgf8a* mRNA revealed that the *fgf8a* 5’UTR contains a Nucleolin consensus binding site – UCCCGA^14^, leading us to hypothesize that Nucleolin post-transcriptionally controls the expression of *fgf8a*. We tested for Nucleolin binding to *fgf8a* mRNA through RNA immunoprecipitation and observed that *fgf8a* mRNA is pulled down with Nucleolin while that of the housekeeping gene *actb* is not (Fig. 5C). This suggests that Nucleolin specifically binds to *fgf8a* mRNA and is required to maintain its stability.

**Figure 5.**
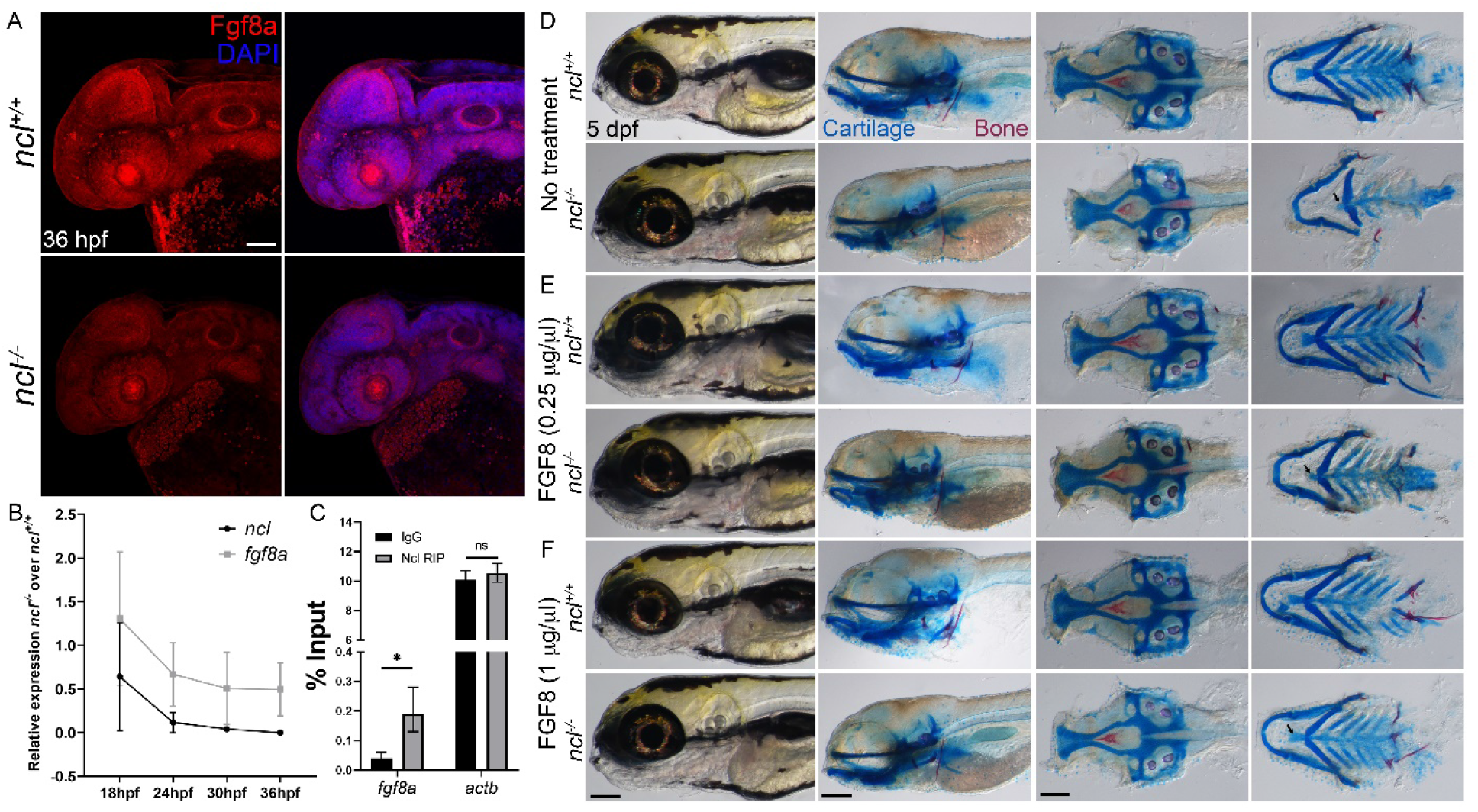
Nucleolin regulates Fgf8a expression. (A) Immunostaining of 36 hpf *ncl^+/+^* and *ncl^-/-^* embryos with an Fgf8a specific antibody reveals reduced expression of Fgf8a in *ncl^-/-^* embryos. (B) qPCR using tissues from *ncl^+/+^* and *ncl^-/-^* embryos at 18, 24, 30 and 36 hpf indicates *ncl* and *fgf8a* expression gradually reduce over time. (C) RNA immunoprecipitation followed by qPCR indicates higher binding of *fgf8a* to Nucleolin compared to IgG control. *actb* was used as negative control. (D) Skeletal preparations of 5dpf *ncl^+/+^* and *ncl^-/-^* larvae as controls for 0.25 μg/μl, (E) and 1μg/μl (F) FGF8 exogenous treatment. Exogenous FGF8 rescues the cranioskeletal phenotype of *ncl^-/-^* larvae. Scale bar denotes 100 μm for A and 70 μm for D, E, and F.

To examine if the phenotype of *ncl^-/-^* mutant embryos is a direct consequence of Fgf8a downregulation, we treated *ncl^+/+^* and *ncl^-/-^* embryos at 18hpf with human recombinant FGF8, which has 76% identity to the zebrafish Fgf8a protein. Compared to untreated *ncl^-/-^* mutant embryos, we observe considerable rescue of the craniofacial cartilage phenotype, which progressively improves with higher concentrations of FGF8 treatment at 60 hpf (Fig. S6 A). The same is true for 5 dpf mutants, with restoration of ceratohyal polarity at 0.25 ug/ul FGF8 (Fig. 5D, E) and rescue of basihyal and ceratobranchial chondrogenesis at 1 ug/ul (Fig. 5D, F). In addition, the length of the trabecula and the parasphenoid is restored in FGF8 treated *ncl^-/-^* larvae. However, while osteogenesis of the tooth improves, the mutant embryos are still missing their 4V^1^ teeth, indicating that FGF8 partially rescues the *ncl^-/-^* phenotype. At 8 dpf, the craniofacial cartilages (Fig. S6 C) as well as the putative posterior swim bladder (Fig. S6 B) are rescued in FGF8 treated *ncl^-/-^* larvae. The FGF8 treated *ncl^-/-^* fry survive at least until 15 dpf, which is 5 days longer than untreated *ncl^-/-^* fry and all the craniofacial skeleton elements are rescued (Fig. S6 D-E). However, these fry are smaller than *ncl^+/+^* fry and their anterior swim bladders fail to inflate. Nonetheless, our data indicates that exogenous FGF8 is sufficient to rescue the craniofacial skeleton defects and increase the lifespan of *ncl^-/-^* mutants.

### FGF8 treatment restores rRNA synthesis in *ncl^-/-^* mutant embryos

*Fgf8* is expressed in the endoderm-derived epithelium of the pharyngeal arches, while Fgf receptors are expressed in both pharyngeal endoderm as well as osteochondroprogenitors^36–38^. Therefore, Fgf8a could directly or indirectly regulate chondrogenic and osteogenic differentiation, suggesting that the FGF8 rescue of cranioskeletal defects of *ncl^-/-^* embryos could be a result of 1) Fgf signaling through FGF8 interaction with Fgf receptor on osteochondroprogenitors, 2) FGF8 could rescue rRNA transcription, or 3) FGF8 could trigger mesenchymal cell-autonomous pathway such as Bmp signaling.

To test if addition of FGF8 rescues rRNA synthesis, we examined rRNA transcription in FGF8 treated *ncl^-/-^* embryos and observe an upregulation in 47S rRNA in FGF8 treated *ncl^-/-^* embryos (Fig. 6A). This could result either by FGF8 directly interacting with rDNA or through Fgfr2, which has been shown to positively regulate rRNA transcription^39^. Consistent with rescued levels of rRNA, the FGF8 treated *ncl^-/-^* embryos also have fewer TUNEL positive cells compared to *ncl^+/+^* embryos (Fig. 6B), indicating that FGF mediated rRNA transcription is important for cell survival.

**Figure 6.**
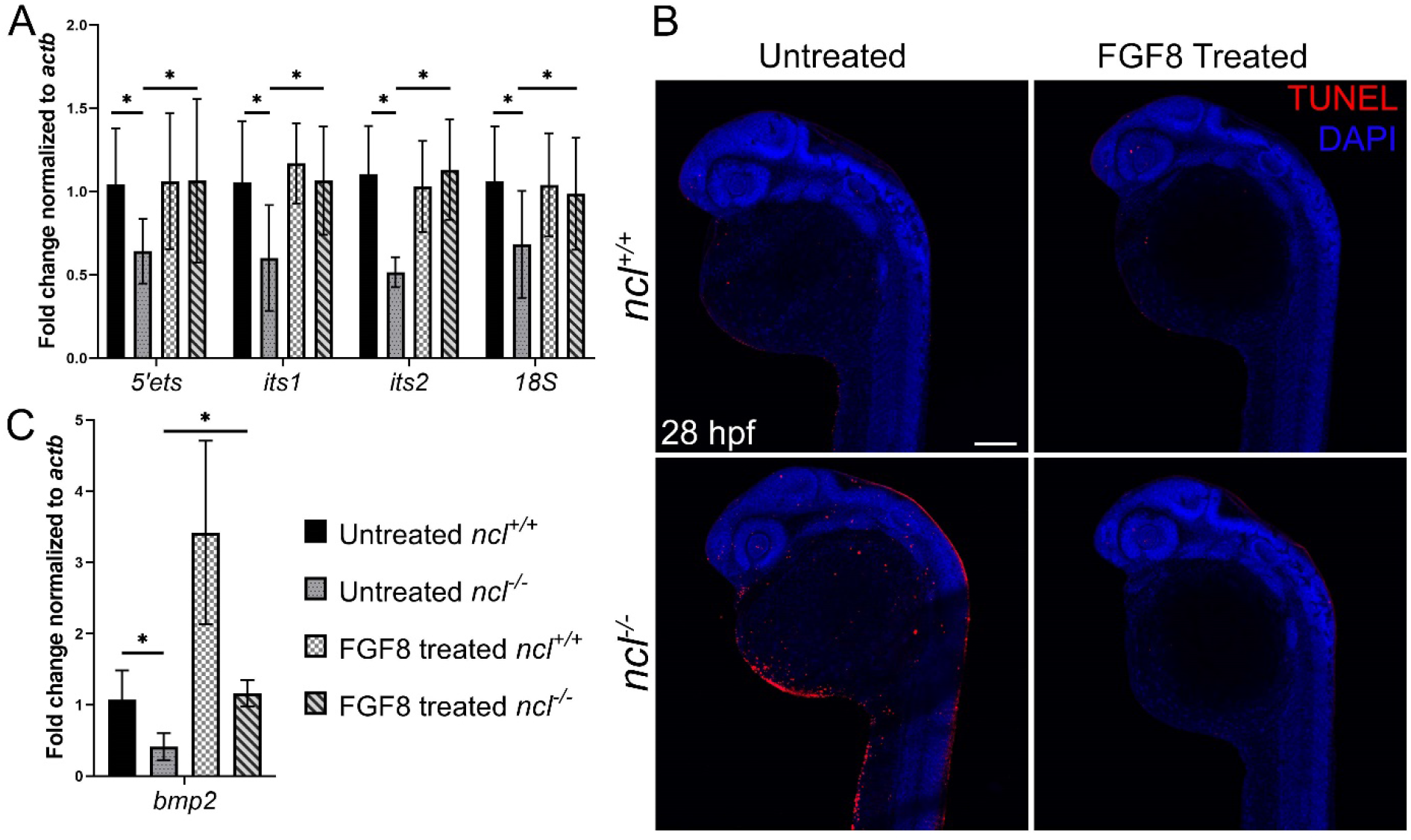
FGF8 rescues rRNA transcription in *ncl^-/-^* embryos. (A) qPCR for *5’ets, its1, its2* and *18S* in untreated and FGF8 treated *ncl^+/+^* and *ncl^-/-^* zebrafish indicates rescue of pre-RNA transcription in FGF8 treated *ncl^-/-^* zebrafish at 36 hpf. (B) TUNEL staining of untreated and FGF8 treated *ncl^+/+^* and *ncl^-/-^* zebrafish at 28 hpf indicates reduced TUNEL positive cells in FGF8 treated *ncl^-/-^* embryos compared to untreated *ncl^-/-^* embryos. (C) qPCR for *bmp2* in 36 hpf *ncl^+/+^* and *ncl^-/-^* embryos untreated and treated with 1 μg/μl FGF8 indicates significant downregulation in *bmp2* in untreated *ncl^-/-^* embryos and significant upregulation in FGF8 treated *ncl^+/+^* embryos. In FGF8 treated *ncl^-/-^* embryos, the *bmp2* transcript level is rescued and comparable to untreated *ncl^+/+^* embryos. *actb* was used as housekeeping control. Scale bar denotes 70 μm for B.

It is well known that Fgf and Bmp signaling interact synergistically during craniofacial development^40^. More specifically, Fgf8 and Bmp2, respectively regulate osteochondroprogenitor differentiation by positively regulating Sox9 expression^41,42^. Therefore, we examined the expression of *bmp2* in FGF8 untreated and treated *ncl^-/-^* embryos at 30 hpf. We observe that while *bmp2* expression is downregulated in untreated *ncl^-/-^* embryos, its expression is increased in treated *ncl^+/+^* and *ncl^-/-^* embryos, suggesting that addition of FGF8 rescues cartilage and bone development by stimulating Bmp signaling in osteochondroprogenitors (Fig. 6C). Taken together, our data suggests that Nucleolin regulates craniofacial development through two different pathways, one by controlling ribosomal RNA transcription and the other by regulating FGF signaling.

## Discussion

Nucleolin is the most abundant phosphoprotein in the nucleolus^43^, however its localization is not limited to the nucleolus. Nucleolin is also observed in the nucleus, cytoplasm and plasma membrane where depending on the cell type and the environmental condition, Nucleolin performs various functions ranging from DNA replication and repair^44^, chromatin remodeling^18^, rRNA transcription and processing^13,20^, mRNA turnover and translation^22,45–51^ and viral entry and replication^52–54^. While the cellular and molecular function of Nucleolin has been previously studied *in vitro*, its role in vertebrate development remains to be explored. Here we show that Nucleolin is critical for zebrafish craniofacial development and specifically for proper differentiation of NCC into cartilage and bone.

Similar to other rRNA modifying proteins such as Nol11, Wrd43 and Fibrillarin whose absence results in chondrogenesis defects in zebrafish and frog^6,11,12^, it is interesting that perturbation of a global process such as ribosome biogenesis results in craniofacial defects. One possible reason for this tissue specificity could be that genes required for rRNA transcription such as subunits of RNA Pol I, *fbl* and *ncl* are highly expressed in NCC and craniofacial tissues, making them more susceptible to disruptions in rRNA transcription^9–11,55^. rRNA transcription is essential not only because it forms the catalytic core of ribosomes, but also because in its absence cells undergo proteotoxic stress as a result of ribosomal protein accumulation^56,57^. Free ribosomal proteins bind to Mdm2 resulting in the accumulation of p53, and consequently cell cycle arrest and cell death^21,58^ (Fig. 7A, B).

**Figure 7.**
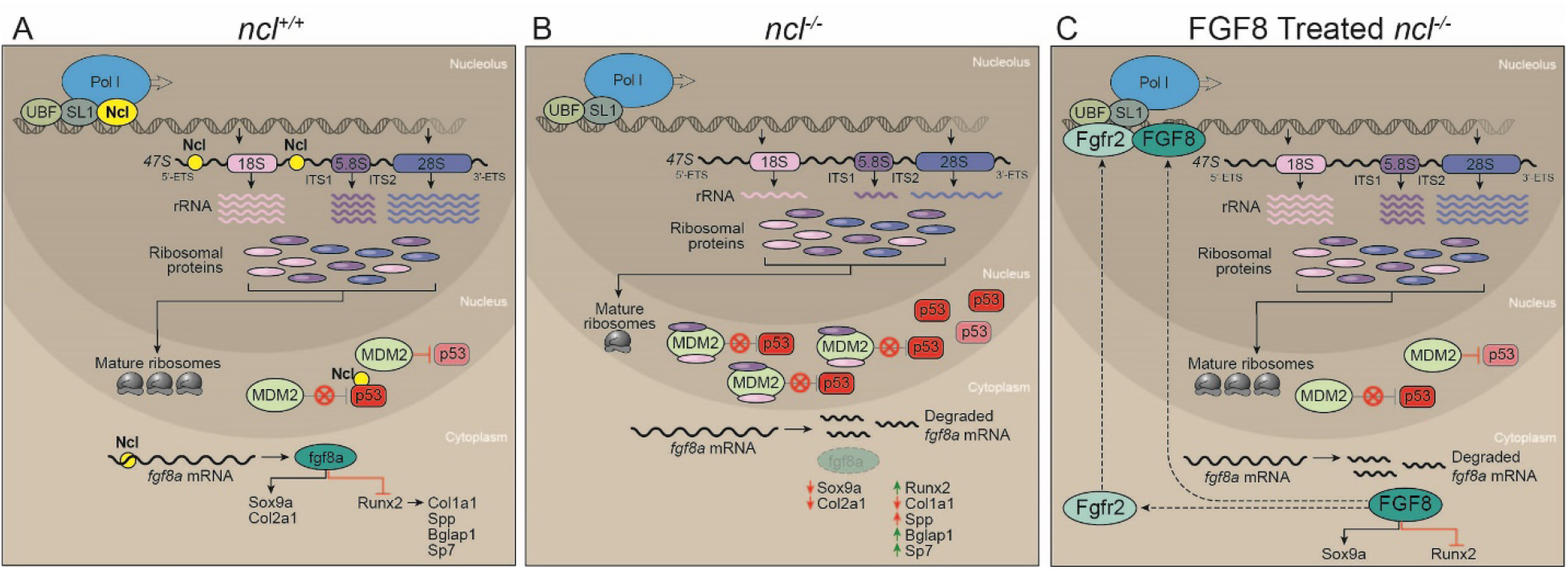
Nucleolin regulates rRNA transcription and fgf8a mRNA stability. (A) In wildtype zebrafish embryos, Nucleolin is required for rRNA transcription as well as processing. The rRNA transcripts are assembled together with ribosomal proteins to make ribosomes. Meanwhile Mdm2 binds to and ubiquitinates p53, which results in p53 proteasomal degradation. Nucleolin binds to and stabilizes p53 protein, thereby acting antagonistically to Mdm2. Furthermore, Nucleolin binds to and stabilizes *fgf8a,* which results in Fgf8a regulated chondrogenesis and osteogenesis, leading to proper craniofacial development. (B) In *ncl^-/-^* embryos, the absence of Nucleolin results in reduced rRNA transcription and possibly, a free pool of ribosomal proteins that bind to Mdm2. This limits Mdm2 binding to p53, resulting in a temporary increase in p53. However, due to the lack of Nucleolin in the cell, p53 protein and *fgf8a* mRNA have reduced half-lives. This results in misregulated chondrogenesis and osteogenesis leading to cranioskeletal defects. (C) In FGF8 treated *ncl^-/-^* embryos, exogenous FGF8 rescues cranioskeletal defects most likely by regulating chondrogenesis and osteogenesis as well recovering rRNA transcription either directly by FGF8 or indirectly by Fgfr2 binding to rDNA.

Contradicting previous *in vitro* studies in human cell lines^59^, we observe that Nucleolin does not bind to *p53* mRNA in zebrafish, most likely due to the differences in the 5’ and 3’ UTR sequences of human and zebrafish p53. However, Nucleolin and p53 proteins bind to each other suggesting that Nucleolin is required to stabilize p53 protein^23^. In the absence of Nucleolin, p53 accumulates temporarily and apoptosis is initiated. However, the effect is not sustained and it does not result in lethality. This is consistent with an initial upregulation of *p21* mRNA expression from 18-24 hpf, followed by a steep decrease by 36 hpf, and is indicative of a momentary surge in the transcriptional activity of p53. p21 is a cell cycle inhibitor, which when significantly reduced releases cell cycle arrest^60,61^, resulting in higher proliferation in *ncl^-/-^* embryos compared to *ncl^+/+^* embryos. This increase in proliferation in the absence of Nucleolin is contrary to published literature, which suggests that Nucleolin promotes proliferation in various cell types^62,63^. This indicates that a different mechanism regulates proliferation in the absence of Nucleolin in mammalian cells and developing zebrafish. Over time, apoptosis decreases, and proliferation increases in *ncl^-/-^* embryos, which results in embryo survival until 5 dpf.

The chondrogenic hypoplasia observed in *ncl^-/-^* mutants can be linked to Sox9a downregulation, while the osteogenic defects arise due to the higher expression of Runx2. This leads to premature osteoblast differentiation, which reduces the pool of cells that can subsequently differentiate into osteoblasts. Thus, early osteoblast markers such as *spp1* and *sp7* are more highly expressed in *ncl^-/-^* embryos compared to *ncl^+/+^* siblings, while late osteoblast markers such as *bglap* are downregulated. However, both Sox9a and Runx2 are indirect targets of Nucleolin based on in silico analysis of their 5’ and 3’ UTRs, which do not contain a Nucleolin consensus binding site. Fgf8a, which is also downregulated in *ncl^-/-^* mutants and is required for osteochondroprogenitor differentiation, does contain a Nucleolin binding site in its 5’ UTR. The 5’ UTR of an mRNA is responsible for both stability and translation of mRNA^64^. Given that *fgf8a* mRNA decreases over time together with reductions in Nucleolin, this suggests that Nucleolin is responsible for stabilizing *fgf8a* mRNA by binding to its 5’ UTR. In addition, the maternal expression of Nucleolin in zebrafish until 18 hpf possibly prevents early embryonic phenotypes such as agenesis of the cerebellum and MHB organizer which have previously been associated with Fgf8a downregulation^65^. This is further corroborated by the restoration of osteogenesis and chondrogenesis in *ncl^-/-^* mutants with exogenous FGF8, suggesting that the critical time for Fgf8a function in *ncl^-/-^* zebrafish is around 18 hpf (Fig. 7C). FGF8 treatment increased the survival of *ncl^-/-^* larvae by 5 days, suggesting continued treatment may further augment lifespan and this will be tested in the future.

In mice Fgf8 has been shown to preferentially bind to Fgfr1^66^. However, in zebrafish Fgf receptors are functionally redundant with respect to Fgf8a ligand binding^67^. Therefore, it is likely that Fgf8a could activate Fgfr2. Fgfr2 is known to bind to the promoter of rDNA and result in histone modification, which activates rRNA transcription similar to Nucleolin^39^. This suggests that upon FGF8 treatment, either FGF8 or Fgfr2 translocates to the nucleolus and changes the state of the rDNA chromatin from closed to open (Fig. 7C). However, given that the upstream core elements of rDNA that RNA Pol I and UBTF bind to are as yet unannotated in zebrafish, further work is required to test our hypothesis. This involves identifying the upstream core elements using a combination of ATAC-seq and ChIP-seq followed by elucidating the mechanism of Fgf8 and Fgfr2 regulation of rRNA transcription in the absence of Nucleolin.

*Fgf8* has been previously shown to regulate variance in facial shape in a dose dependent manner^68^. Furthermore, rDNA transcription is known to be essential for craniofacial development^69,70^ and variation^71^. Therefore, as an upstream regulator of *Fgf8,* and with a role in rDNA transcription, by extrapolation, Nucleolin may also play an important role in determining facial shape. Overall, our work uncovered that Nucleolin regulates Fgf8 signaling as well as rRNA transcription, making craniofacial development especially susceptible to Nucleolin loss-of-function.

## Materials and Methods

### Zebrafish

Adult zebrafish (Danio rerio) were housed and maintained in the Stowers Institute Zebrafish Facility according to an IACUC approved protocol (# 2021-124). Zebrafish embryos were raised at 28.5°C and staged using standard procedures^72^. To prevent pigment development for immunostaining experiments 0.002% 1-Phenyl-2-thiourea was added to the embryo media. *ncl^hi2078Tg^* zebrafish were obtained through ZIRC and maintained as heterozygotes on the AB/TU background and incrossed to generate homozygous mutant embryos. PCR was used to detect for the presence or absence of the insertional mutation. The *ncl* wild type allele was detected using the following primers: forward 5’-TTACATGTGGTGAGAAGGCCC-3’ and reverse 5’-AACACCTCCCCTGGGTTTAT-3’. The *ncl* mutant allele was detected using the following primers: forward 5’-TTACATGTGGTGAGAAGGCCC-3’ and reverse 5’-GCTAGCTTGCCAAACCTACAGGT-3’. The *ncl* heterozygous mutant lines were crossed with reporter line *Tg(7.2kb-sox10:gfp)*, referred to as *sox10:egfp*, as well as the *tp53^M214K^* line.

### Live imaging

Embryos were anaesthetized with MS-222 and mounted in 2% methyl cellulose while submerged in E2 media. Embryos were imaged using a Leica MZ16 microscope equipped with a Nikon DS-Ri1 camera and NIS Elements BR 3.2 imaging software. When appropriate, manual Z stacks were taken and the images were assembled using Helicon Focus software.

### Skeletal stain

Alcian blue and Alizarin red staining to label cartilage and bone, respectively, was performed according to Walker and Kimmel^72^. Embryos were cleared in glycerol and potassium hydroxide and dissected for neurocranium and viscerocranium images. Imaging was performed as described above.

### Immunostaining, EdU and TUNEL

Whole-mount immunostaining was performed according to standard protocols^73^ using primary antibodies against Nucleolin (1:500, abcam #ab22758), GFP (1:500, Life Technologies #A6455), Zn-8 (1:250, DSHB), Sox10 (1:500, GeneTex # GTX128374), Sox9a (1:500, GeneTex # GTX128370), Runx2 (1:500, Abcam # ab23981), Fgf8a (1:500, GeneTex # GTX128126) and pHH3 (1:2000, Millipore # 06-570). Fluorescent secondary antibodies, either Alexa-488 or Alexa-546 (1:500, Invitrogen) were used for detection. TUNEL and EdU Click-IT assay were performed according to the manufacturer’s instructions with slight modifications. Embryos were incubated for 1 hour on ice and 1 hour at 37°C in the reaction buffer. Embryos were imaged using a Zeiss upright 700 confocal microscope and images were captured and processed using Zen software. ImageJ software was used to quantify the fluorescence intensity and area. The Student’s t-test was used to determine statistical significance.

### Alkaline Phosphatase staining

Fixed embryos were washed in Tris-Buffer Saline and NTMT buffer (100mM NaCl, 100mM Tris-HCl, pH9.5, 50mM MgCl2, 1% Tween20), followed by incubation with NBT (3.5µl) and BCIP (5µl) in NTMT for the desired color revelation time. The images were collected using a Nikon DS-Ri1 camera.

### Western blot

Protein samples consisting of 5 fish/sample were collected at appropriate stages. Embryos were homogenized and suspended in sample buffer containing Tris pH 8.0, Sodium Chloride, SDS, Sodium deoxycholate, NP-40 and protease inhibitor and used for Western blotting according to standard protocols^74^. Protein quantity was estimated via a BCA assay. Primary antibodies used were γ-tubulin (1:1000, Sigma), α-tubulin (1:10000, Sigma), p53 (1:500, Cell Signaling Technology, #2524S) and Nucleolin (1:500, Abcam #ab22758). Western blots were imaged and quantified using a CLx-Scanner (Li-COR) and Odyssey Software. For quantification, band intensities for p53 were compared to housekeeping control ɣ-Tubulin. The Student’s t-test was performed for statistical analysis.

### Immunoprecipitation

Protein lysates from 28 hpf control and *ncl^-/-^* zebrafish were used (n = 25 per biological replicate; total three biological replicates) for pre-conjugation with Mdm2 antibody (Cell Signaling Technology, #86934) and IgG with magnetic beads. Because the embryos were group and lysed immediately after collection, genotyping by conventional methods could not be performed. Instead, the lysate was used in a western blot for Nucleolin to substitute as genotyping. Following immunoprecipitation, the samples were used in western blotting and immunoblotted for both Mdm2 and p53.

### qPCR

RNA was collected from zebrafish embryos using the Qiagen miRNeasy Micro Kit and tested for quality on an Agilent 2100 Bioanalyzer. The Superscript III kit (Invitrogen) was used to synthesize cDNA for qPCR using random hexamer primers. The following primers were used: ncl forward 5’-ATATCGAGGGCAGGAGTATT-3’ and reverse 5’-GTTTTCGTAGGTCCAGAGTT-3’; tp53 forward 5’-CGAGCCACTGCCATCTATAAG-3’ and reverse 5’-TGCCCTCCACTCTTATCAAATG-3’, p21 forward 5’-GACCAACATCACAGATTTCTAC-3’ and reverse 5’-TGTCAATAACGCTGCTACG-3’, sox9a forward 5’-GGAGCTCAGCAAAACTCTGG-3’ and reverse 5’-AGTCGGGGTGATCTTTCTTG-3’; col2a1 forward 5’-GCGACTTTCACCCCTTAGGA-3’ and reverse 5’-TGCATACTGCTGGCCATCTT-3’; runx2a forward 5’-AACTTTCTGTGCTCGGTGCT-3’ and reverse 5’-AACTTTCTGTGCTCGGTGCT-3’; runx2b forward 5’-CAAACACCCAGACCCTCACT-3’ and reverse 5’-GTATGACCATGGTGGGGAAG-3’; col1a2 forward 5’-CTGGCATGAAGGGACACAG-3’ and reverse 5’-GGGGTTCCATTTGATCCAG-3’; col10a1 forward 5’-CCTGTCTGGCTCATACCACA-3’ and reverse 5’-AAGGCCACCAGGAGAAGAAG-3’; fgf8a forward 5’-GCCGTAGACTAATCCGGACC-3’ and reverse 5’-TTGTTGGCCAGAACTTGCAC-3’; osteocalcin forward 5’-TGAGTGCTGCAGAATCTCCTAA-3’ and reverse 5’-GTCAGGTCTCCAGGTGCAGT-3’; osteopontin forward 5’-TGAAACAGATGAGAAGGAAGAGG-3’ and reverse 5’-GGGTAGCCCAAACTGTCTCC-3’; sp7 forward 5’-GGATACGCCGCTGGGTCTA-3’ and reverse 5’-TCCTGACAATTCGGGCAATC-3’; bmp2 forward 5’-TCCATCACGAAGAAGCCGTGG-3’ and reverse 5’-TGAGAAACTCGTCACTGGGGA-3’; actb forward 5’-TTCCTTCCTGGGTATGGAATC-3’ and reverse 5’-GCACTGTGTTGGCATACAGG-3’; canx forward 5’-ACGATACCGCAGAGAATGGAGACA-3’ and reverse 5’-TCCTGTTTCTGGGAGACCTCCTCA-3’. Previously published rRNA primer sequences were used for qPCR^75^. Power Sybr (Life Technologies) reaction mix and the ABI 7900HT real time PCR cycler was used measure cDNA amplification. Three biological replicates were run in technical triplicate for each experiment. No template and no reverse-transcriptase samples were run as negative controls.

### RNA immunoprecipitation

Lysates from 28hpf wild-type zebrafish were used (n = 100 per biological replicate; total three biological replicates) for pre-conjugation with Nucleolin antibody (Abcam, # ab22758) and IgG with magnetic beads. Immunoprecipitation was performed using manufacturer’s instructions for the EZ-Nuclear RIP Kit (EMD Millipore, #17-10 523), followed by cDNA synthesis and qRT-PCR for 5’ETS, ITS1, ITS2 and 18S rRNA, *fgf8a*, *p53* and *actb*.

### FGF8 treatment

*ncl^+/+^* and *ncl^-/-^* embryos were treated with 0.0625, 0.25 and 1 ng/ul human recombinant FGF8 (Thermofisher scientific, # PHG0184) diluted in E2 media by immersion starting at 18hpf. The FGF8 in media was replaced every 24 hours until 5 dpf following which the larvae were added to the tank system and allowed to develop until 15 dpf.

### Morpholino knockdown

1-2 cell stage *ncl^+/+^* and *ncl^-/-^* embryos were injected with 5–10 ng of morpholino (MO) targeting the p53 AUG start site or a control MO (Gene Tools, LLC). The sequences of the MOs used were as follows: p53 MO 5’-GCGCCATTGCTTTGCAAGAATTG-3’ and control MO 5’-CCTCTTACCTCAGTTACAATTTATA-3’. Following injection, embryos were raised at 28°C to 5 dpf for skeletal staining.

## Acknowledgements

We are thankful to all members of the Trainor lab for their input and suggestions during the course of this work. We thank Dr. Tom Schilling for the gift of the *sox10:egfp* zebrafish, Dr. Tatjana Piotrowski for the gift of the tp53 zebrafish and the SIMR Aquatics staff, especially Carrie Carmichael for zebrafish care and maintenance. We also thank Mark Miller for illustrating Figure 7 and Dr. Cathy McKinney for imaging support. This work was supported by the Stowers Institute for Medical Research (P.A.T) and a postdoctoral fellowship from American Association for Anatomy (S.D). Original data underlying this manuscript can be accessed from the Stowers Original Data Repository at http://www.stowers.org/research/publications/libpb-1658.

---

**Figure S1.**
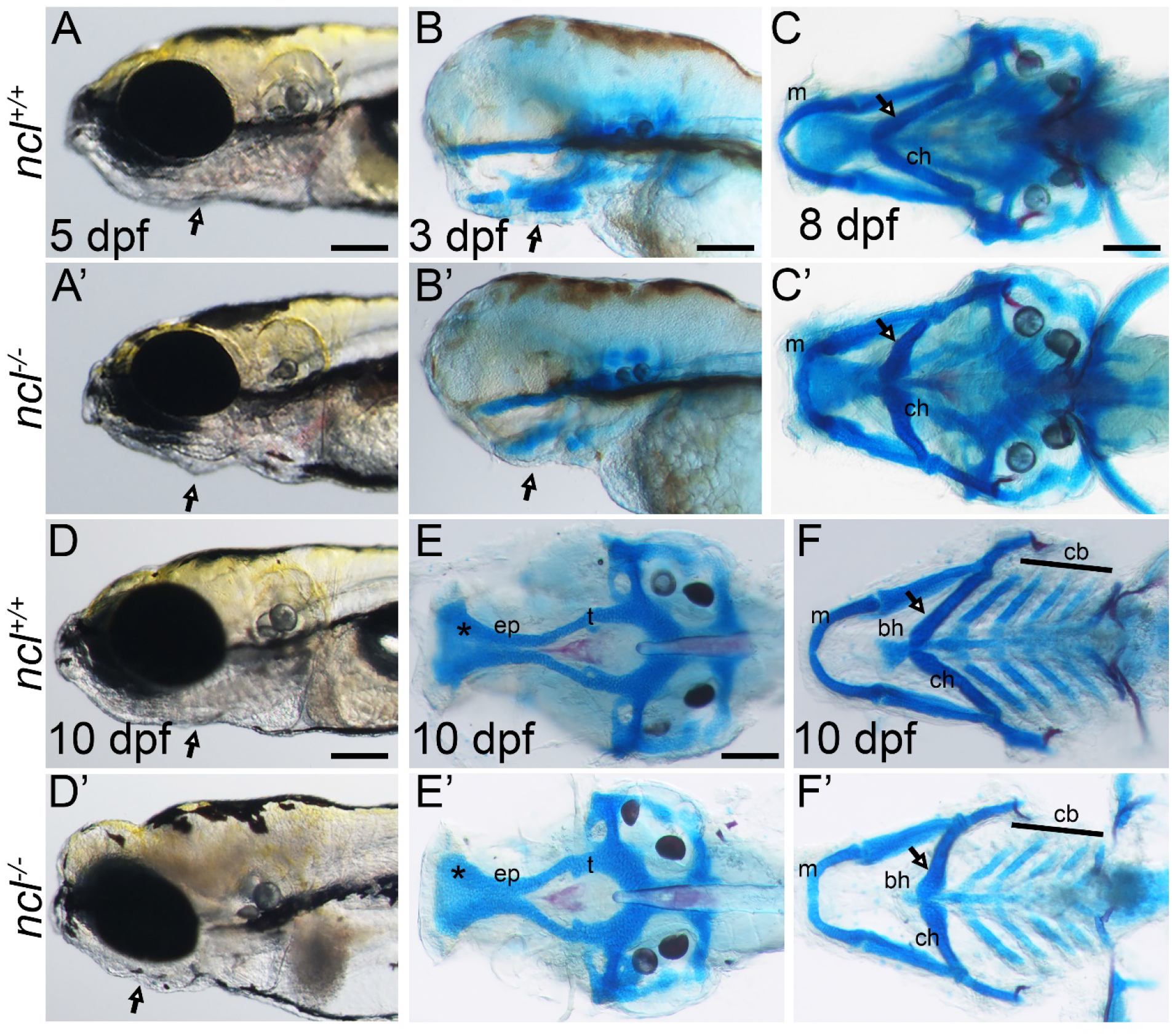
*ncl^-/-^* mutants exhibit craniofacial defects and embryonic lethality. (A-A’) Brightfield imagining of 5 dpf *ncl^+/+^* and *ncl^-/-^* reveals the lower jaw is smaller in *ncl^-/-^* larvae (black arrow). Skeletal staining of 3 dpf (B-B’) and 8 dpf (C-C’) *ncl^+/+^* and *ncl^-/-^* larvae shows the ceratohyal in hypoplastic in 3 dpf mutants (B’) and misshapen in 8 dpf mutants (C’). (D-D’) Brightfield imagining of 10 dpf *ncl^+/+^* and *ncl^-/-^* larvae indicates the lower jaw is smaller in *ncl^-/-^* larvae (black arrow) and reveals necrotic tissue in the craniofacial region and the heart. Skeletal staining with Alcian blue shows differential labeling of the ethmoid plate (asterisk), with smaller trabecula in the neurocranium in 10 dpf *ncl^-/-^* larvae compared to controls (E-E’). Furthermore, in the viscerocranium, Meckel’s cartilage is bent, the basihyal and ceratobranchials are hypoplastic and the angle of the ceratohyal is obtuse (F-F’). Abbr. ep: ethmoid plate, t: trabecula, m: meckel’s, bh: basihyal, ch: ceratohyal, cb: ceratobranchial. Scale bars denotes 350 μm for A-A’, 50 μm for B-B’, 100 μm for C-F’.

**Figure S2.**
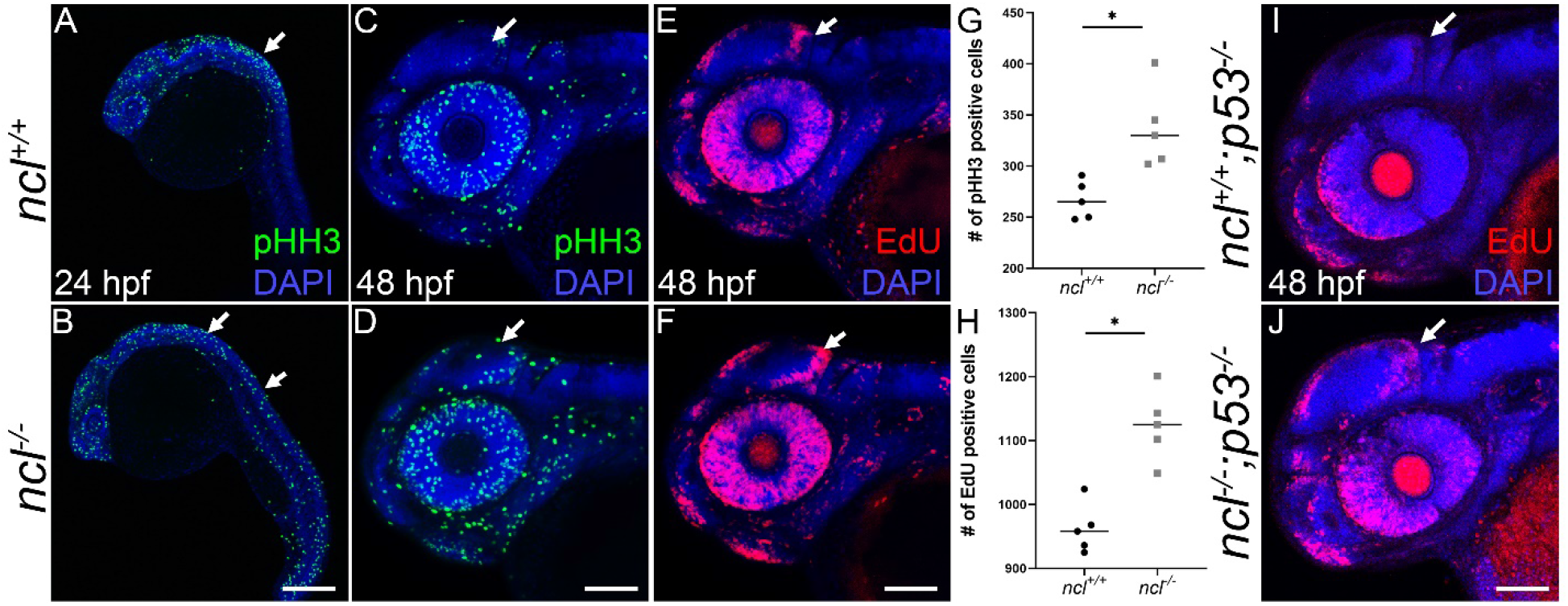
Proliferation is increased in *ncl^-/-^* embryos. At 24 hpf, proliferating cells were labeled with the mitotic marker, pHH3, which revealed no significant difference between *ncl^+/+^* (A) and *ncl^-/-^* (B) embryos. By 48 hpf, the number of proliferating cells labeled with pHH3 (C-D) and EdU (E-F) is significantly higher in the *ncl^-/-^* embryos compared to *ncl^+/+^* embryos, especially in the midbrain-hindbrain boundary. Quantification of pHH3 (G) and EdU (H) positive cells in the *ncl^+/+^* and *ncl^-/-^* embryos. Similarly, *ncl^-/-;^p53^-/-^* embryos (J) have more proliferating cells than *ncl^+/+^* embryos (I). Scale bar denotes 70 μm for A-F, I-J.

**Figure S3.**
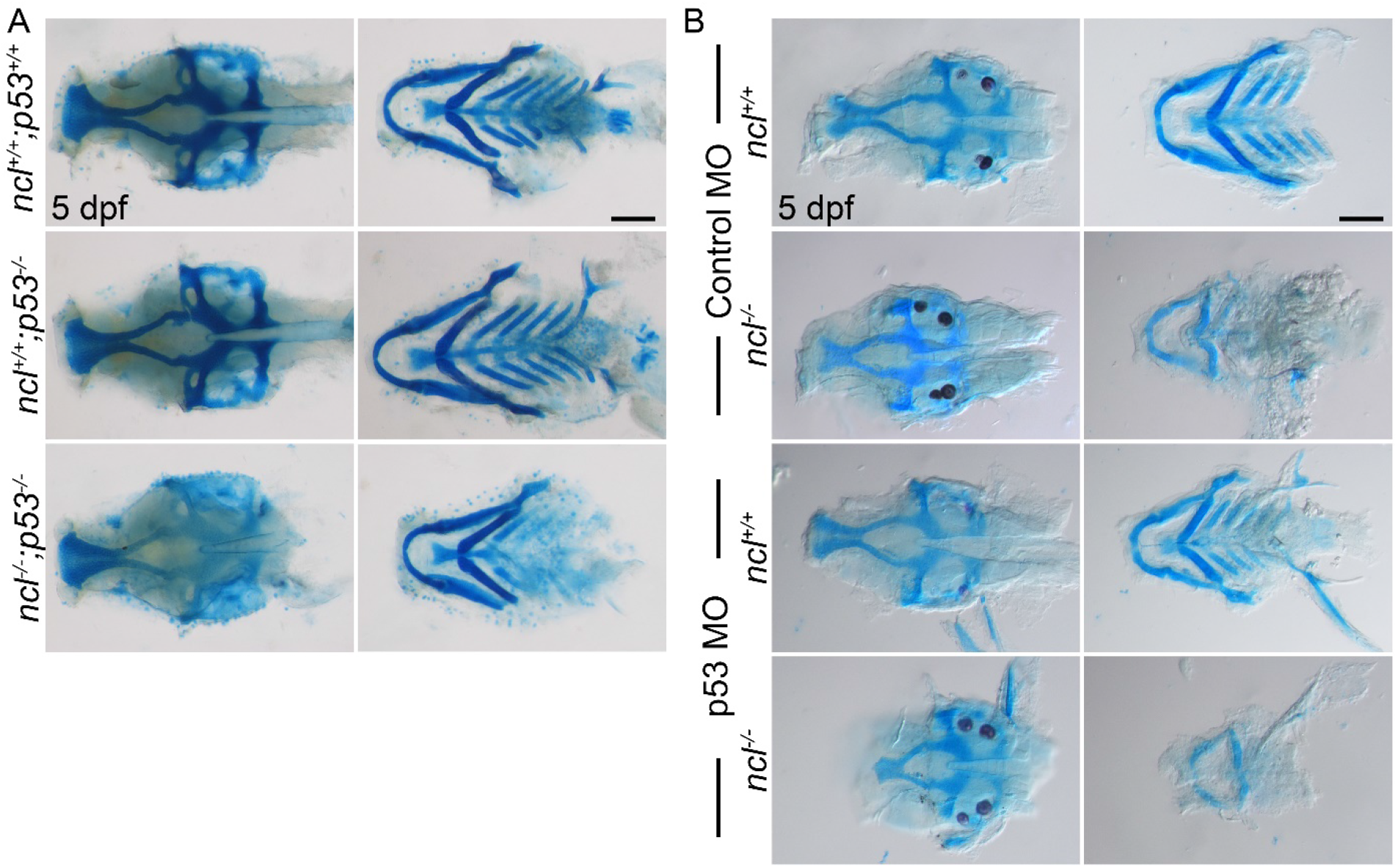
p53 downregulation does not rescue the cranioskeletal anomalies in *ncl^-/-^* mutants. (A) Compared to *ncl^+/+^;p53^+/+^* and *ncl^+/+^;p53^-/-^* larvae at 5 dpf, *ncl^-/-^;p53^-/-^* have hypoplastic neurocranium and ceratohyal cartilages. (B) Microinjecting p53 morpholinos into one cell stage *ncl^+/+^* and *ncl^-/-^* embryos reduces the size of the neurocranium and ethmoid plate as well as chondrogenesis of the ceratobranchials in *ncl^-/-^* larvae compared to control morpholino injected *ncl^-/-^* larvae. Abbr. MO: morpholino. Scale bar denotes 70 μm for A-B.

**Fig S4.**
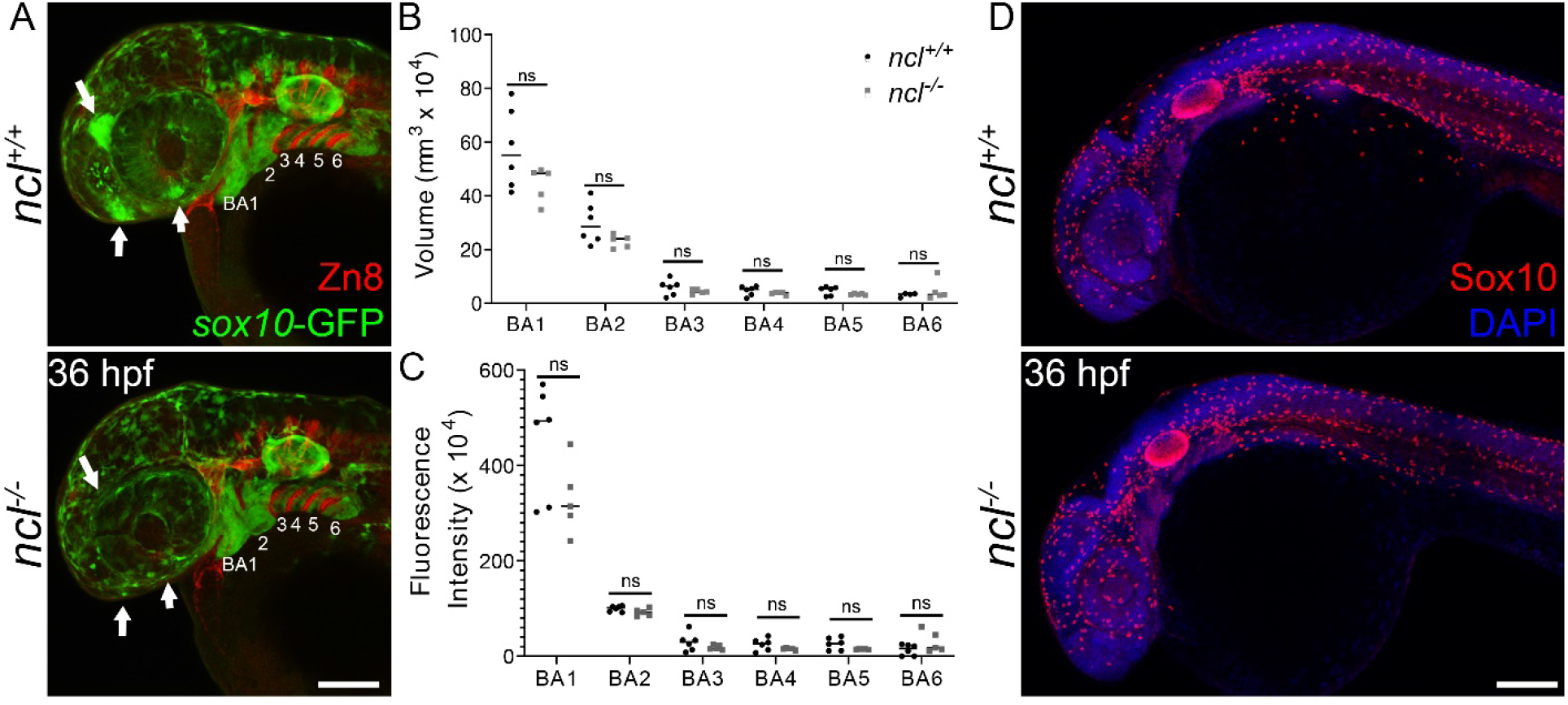
Neural crest cell induction and migration is unaffected in *ncl^-/-^* mutant embryos. (A) *ncl^+/+^* and *ncl^-/-^* embryos at 36 hpf were immunostained with YFP to reveal NCC labeled by the *sox10-egfp* transgene, and for the pharyngeal pouch marker, Zn8 to demarcate each pharyngeal arch. Volumetric (B) and fluorescence intensity (C) analysis of the pharyngeal arches indicates that the arches in *ncl^+/+^* and *ncl^-/-^* embryos are of comparable volume and have similar amounts of NCC. (D) Sox10 immunostaining reveals similar numbers of Sox10 positive cells in *ncl^+/+^* and *ncl^-/-^* embryos. Scale bar denotes 100 μm for A and D.

**Fig. S5.**
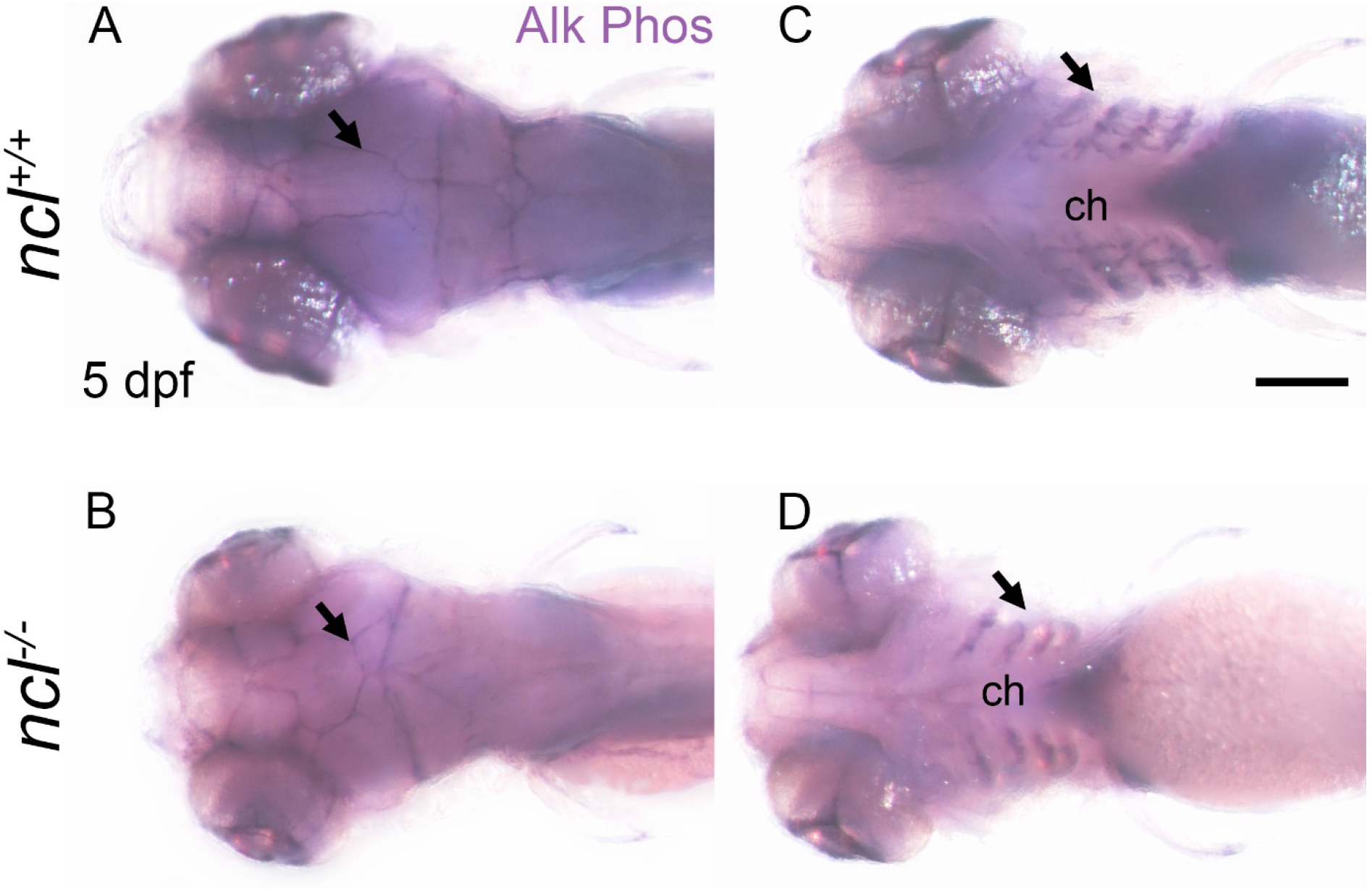
Osteoblast population is reduced in *ncl^-/-^* embryos. Alkaline phosphatase staining reveals that compared to 5 dpf *ncl^+/+^* embryos (A), the sutures are misshapen in 5 *ncl^-/-^* embryos (B). Similarly, in ventral view, compared to *ncl^+/+^* embryos (C), the intensity of alkaline phosphatase staining is significantly reduced in the pharyngeal arches in *ncl^-/-^* embryos (D). Scale bar denotes 140 μm for A and D.

**Fig S6.**
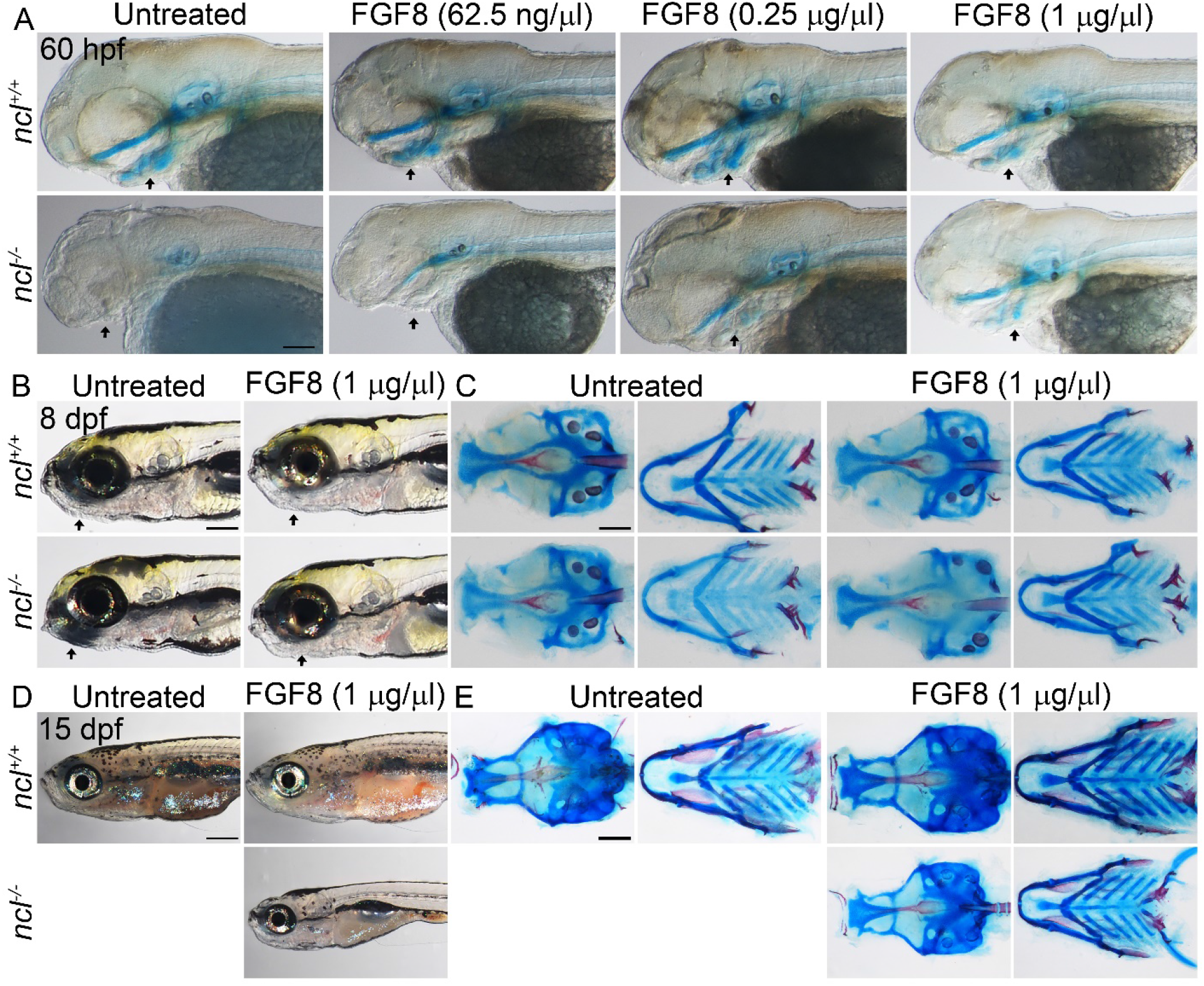
FGF8 treatment rescues *ncl^-/-^* embryos. (A) Alcian blue and Alizarin red staining of *ncl^+/+^* and *ncl^-/-^* embryos treated with FGF8 indicates that with increasing dose of FGF8, the cartilage elements of 60hpf *ncl^-/-^* embryos are progressively rescued. (B) Brightfield images of 8 dpf *ncl^+/+^* and *ncl^-/-^* larvae show that with FGF8 treatment, the craniofacial and swim bladder phenotypes are rescued. (C) Skeletal staining of 8 dpf untreated and treated *ncl^+/+^* and *ncl^-/-^* larvae indicates that most of the cranioskeletal defects including the hypoplastic basihyal and cetatobranchial are rescued with 1 μg/μl FGF8 treatment. (D) Brightfield images of 15 dpf *ncl^+/+^* and *ncl^-/-^* larvae show that with FGF8 treatment the larvae are smaller compared to untreated and treated *ncl^+/+^* larvae. In addition, the anterior swim bladder is not inflated in the treated *ncl^-/-^* larvae. The untreated *ncl^-/-^* larvae do not survive beyond 10 dpf. (E) Skeletal staining of 15 dpf untreated and treated *ncl^+/+^* and *ncl^-/-^* larvae indicates that most of the cranioskeletal defects are comparable between treated *ncl^+/+^* and *ncl^-/-^* larvae, although the embryo is smaller. Scale bar denotes 50 μm for A, 250 μm for B, 100 μm for C, 350 μm for D and 200 μm for E.

